# A network of CLAVATA receptors buffers auxin-dependent meristem maintenance

**DOI:** 10.1101/2022.05.27.493758

**Authors:** Amala John, Elizabeth Sarkel Smith, Daniel S. Jones, Cara L. Soyars, Zachary L. Nimchuk

**Affiliations:** Department of Biology, University of North Carolina at Chapel Hill, 27599 Chapel Hill, NC, USA; Curriculum in Genetics and Molecular Biology, University of North Carolina at Chapel Hill, 27599 Chapel Hill, NC, USA

## Abstract

Plant body plans are elaborated in response to both environmental and endogenous cues. How these inputs intersect to promote growth and development remains poorly understood. During reproductive development, central zone stem cell proliferation in inflorescence meristems is negatively regulated by the CLAVATA3 (CLV3) peptide signaling pathway. In contrast, floral primordia formation on meristem flanks requires the hormone auxin. Here we show that CLV3 signaling is also necessary for auxin-dependent floral primordia generation and that this function is partially masked by both inflorescence fasciation and heat-induced auxin biosynthesis. Stem cell regulation by CLAVATA signaling is separable from primordia formation but is also sensitized to temperature and auxin levels. Additionally, we uncover a novel role for the CLV3 receptor CLAVATA1 in auxin-dependent meristem maintenance in cooler environments. As such, CLV3 signaling buffers multiple auxin-dependent shoot processes across divergent thermal environments with opposing effects on cell proliferation in different meristem regions.

## Main

Plant development is highly tuned to environmental conditions such as light, daylength, and temperature^1^. Despite this, components of the plant body plan, like inflorescences and flowers, develop robustly across diverse environmental conditions^2^. How core plant developmental mechanisms accommodate environmental fluctuations to maintain developmental robustness remains poorly understood and impacts the ability to create climate change resilient crops^3^.

Inflorescence formation and flower production in Arabidopsis are robust across thermal environments^2^. Flower formation on inflorescence meristems (IM) requires the hormone auxin^4, 5^. Flowers initiate as primordia on IM flanks where auxin maxima form before proliferating and differentiating into floral meristems (FMs) that subsequently generate floral organs^6^. Mutations that abolish auxin function give rise to classical “pin-formed” shoots lacking floral primordia^7^. Continuous flower production and shoot growth is supported by balanced stem cell proliferation in the IM center. Stem cell proliferation is negatively regulated by CLAVATA3, a founding member of the CLE peptide (CLEp) family of extracellular ligands, of which there are 32 in Arabidopsis^8, 9^. Mutations that reduce CLV3p signaling cause IM expansion, stem enlargement and widening (fasciation), increases in floral organs such as carpels, and are also associated with yield traits in crop species^10, 11^. CLV3p is produced by central zone stem cells and diffuses into the underlying meristem organizing center (OC) where it signals via the Leucine Rich Repeat (LRR) receptor CLV1 and a dimer consisting of the LRR receptor-like protein CLV2 and the transmembrane pseudokinase CORYNE (CRN)^12–16^. Collectively CLV3p receptors signal to dampen the expression of *WUSCHEL* (*WUS*)^17, 18^, which encodes a stem cell promoting transcription factor^19^. Additionally, CLV1 signaling also represses the expression of three related *BARELY ANY MERISTEM* (*BAM1-3*^20^) LRR receptor kinases in the OC as part of a *WUS*- independent transcriptional compensation mechanism buffering the loss of *CLV1*^21, 22^. Transcriptional compensation is a conserved feature in *CLV* stem cell networks but evolved different wiring across species. For instance, in tomato loss of *CLV3* is partially compensated by transcriptional upregulation of *CLE9*^23^. In Arabidopsis, stem cell defects in *clv1*/*bam* quadruple mutants are considerably stronger than *clv3* mutants, suggesting redundancy between *CLV3* and other *CLE* genes^21, 22^.

Despite being studied for almost 30 years, the relationship between CLV1/BAM and CLV2/CRN receptors in CLEp signaling remains unresolved. CLV2/CRN are additive with CLV1/BAM in stem cell regulation and are not required for *CLV1*/*BAM* transcriptional compensation^14, 22^ or CLV1 endomembrane trafficking in response to CLV3p^24^. CLV1/BAM ectodomains bind CLE peptides^25–29^, but CLV2 appears to lack CLEp binding capability^30^. *CLV2*/*CRN* IM expression differs from *CLV1*^21, 31, 32^, with higher levels in developing primordia and OC cells respectively, despite overlap in other IM regions. Additionally, some *CLV1/BAM* processes outside of shoots do not appear to require *CLV2*/*CRN*^25, 33^. However, CLV1/BAM and CLV2/CRN both require CLV3 INSENSITIVE KINASE (CIK) family co-receptor LRR-kinases to signal^31, 34–37^.

Recently we discovered that CLV2/CRN signaling promotes auxin-dependent floral primordia outgrowth^31^, likely by promoting the expression of *YUCCA* (*YUC*) genes involved in auxin biosynthesis^38, 39^. Upon flowering the primary shoot in *clv2/crn* plants will produce 1-5 normal flowers then stall stem elongation while continuing to initiate primordia that terminate outgrowth around stage 2 prior to becoming FMs. After this termination phase, normal shoot elongation and flower production recovers for reasons unknown (Extended Data Fig. 1h). *CLV2*/*CRN* function in floral primordia generation is only evident in ambient/cooler temperatures as it is masked by *YUC*- dependent auxin biosynthesis from the thermomorphogenesis response pathway in warmer environments. Consistent with this, mutations in the thermomorphogenesis repressor *EARLY FLOWERING3* (*ELF3*^40, 41^) restore primordia formation and shoot elongation to *crn* mutants in cooler environments^31^. The function of CLV2/CRN in stem cell regulation and floral primordia outgrowth appears genetically separable and aborted primordia in *crn* plants are not associated with *WUS* expression^31^. Whether other auxin-dependent IM functions are affected by CLEp signaling, or if other CLEp developmental processes are buffered by temperature, is unclear. Additionally, it is unclear which CLE peptides promote primordia formation or if CLV/BAM receptors participate in primordia formation as existing *cle* and *clv1*/*bam* receptor mutants do not display *clv2/crn*-like primordia defects.

**Fig. 1:**
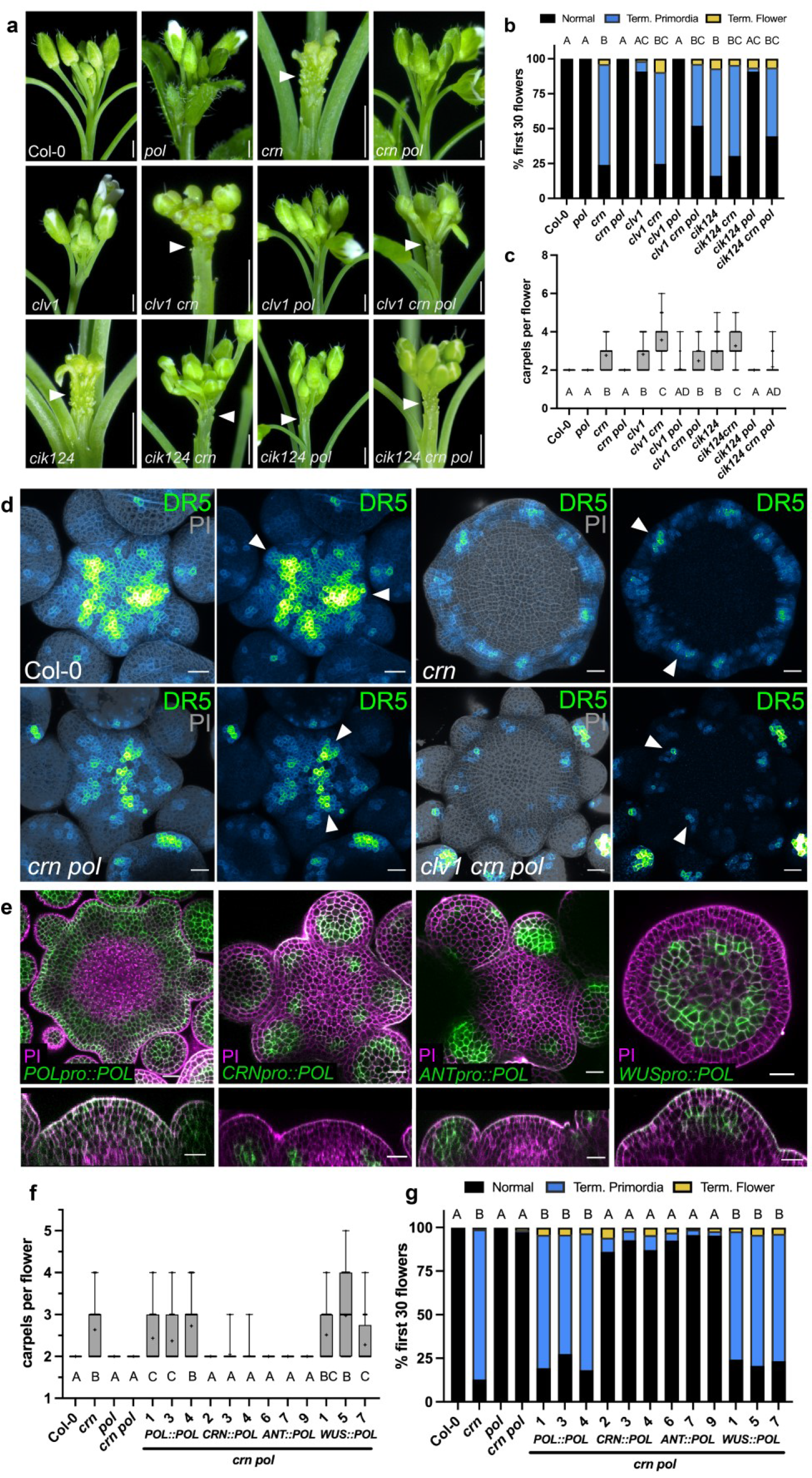
Dual CLV3p receptor systems regulate auxin-mediated primordia outgrowth. a-d, CLV1 and CLV2/CRN signaling promote auxin dependent floral primordia formation. **a,** Inflorescences of Col-0, *pol*, *crn*, *crn pol, clv1*, *clv1 crn*, *clv1 pol*, *clv1 crn pol*, *cik124*, *cik124 crn*, *cik124 pol*, *cik124 crn pol*. Note different magnifications used to highlight primordia termination (arrow). **b**, Quantification of flower primordia termination by categorizing the first 30 attempts to make a flower as normal (black), terminated primordia (blue) or terminated flower (yellow) and **c**, quantification of carpels per flower for Col-0 (n=16), *pol* (n=16), *crn* (n=12), *crn pol* (n=24), *clv1- 101* (n=10), *clv1-101 crn* (n=10), *clv1-101 pol* (n=17), *clv1-101 crn pol* (n=14), *cik124* (n=12), *cik124 crn* (n=12), *cik124 pol* (n=24), *cik124 crn pol* (n=24). All *cik* alleles in this figure are *cik1- 3, cik2-3, and cik4-3* CRISPR generated alleles. **d**, Maximum intensity projection of DR5::GFP (green fire blue LUT) with propidium iodide (PI, grey) and DR5::GFP alone in the IMs of Col-0 (n=9), *crn* (n=9), *crn pol* (n=7), *clv1-101 crn pol* (n=8) grown at 16°C. Arrows point to presumptive incipient P0 and P-1 primordia. (**e-g)** POL regulates CLV1 mediated primordia formation and stem cell regulation from OC cells in *crn pol*. **e**, Confocal imaging with PI staining (magenta) shows expression in XY and Z-axis IM images from *POL::POL-YPET, CRN::POL- YPET, ANT::POL-YPET*, and *WUS::POL-YPET* (green, n=4). **f**, Quantification of carpels per flower and **g**, quantification of flower primordia termination for Col-0, *crn*, *pol*, *crn pol* and *crn pol* transformed with *POL::POL-YPET, CRN::POL-YPET, ANT::POL-YPET,* and *WUS::POL-YPET*. Numbers indicate individual transformants quantified in T2 generation with consistent results (n=15-25). Statistical groupings based on significant differences using Kruskal-Wallis and Dunn’s multiple comparison test correction (**b, c, f, g**) where significance is defined as p-value <0.05 and “+” indicates mean. Scale bars 10mm (**a**) and 20µm (**d, e**).

### Auxin-dependent primordia formation requires overlapping CLV receptor pathways

Previously we noticed that mutations in the membrane associated protein phosphatase *POLTERGEIST* (*POL*, *pol-6*^42, 43^) restored primordia formation in *crn* null mutant plants (*crn-10*) grown in ambient temperatures^31^, but the basis for this was unclear. *pol* mutations are known to repress stem cell defects in diverse *clv* class mutants to varying degrees^43, 44^. We recently discovered that POL dampens CLEp signaling by dephosphorylating CLV1/BAM class receptors, providing a mechanistic explanation for POL function^45^. Therefore, we asked if de-repressed CLV1 signaling was responsible for the restoration of primordia outgrowth in *crn pol* plants. To quantify primordia formation^31^, we grew plants in cool (16-17°C) and ambient temperatures (22- 23°C) and scored the first 30 flower attempts as either normal (fully formed flowers with all organ whorls present), terminated flower primordia (no flower organs develop), or terminated flowers (some flower organs develop, but no gynoecium). Indeed, the null *clv1-101* mutation significantly reduced primordia outgrowth in *crn pol* plants, with stronger effects in cool temperatures (Fig. 1a,b and Extended Data Fig. 1a,b). We confirmed this result with an independent *clv1* allele, the strong *clv1-8* alelle^12^, thought to abolish CLV3p/CLEp binding^29^ (Extended Data Fig. 1c,d). As such, CLV1-promotes floral primordia formation in *crn pol* plants, despite playing a minimal role in *CRN POL* backgrounds. In contrast, *cik1*/*2*/*4* triple mutants displayed significant reductions in primordia formation in both *CRN POL* and *crn pol* backgrounds (Fig. 1a,b), consistent with CIKs mediating both CLV2/CRN and CLV1 signaling^36^. *CIK* genes contributed redundantly to primordia formation and were variably expressed in incipient primordia and OC cells (Extended Data Fig. 2a-c). Primordia outgrowth defects did not track exclusively with increased carpel numbers in flowers, a measure of FM stem cell proliferation, across *clv*/*cik* genotypes demonstrating that these processes are genetically separable (Fig. 1b,c). In all mutant backgrounds, primordia outgrowth was variably restored in warmer temperatures (30^°^C), with lower restoration rates in stronger *clv* mutants (Extended Data Figs. 1a-d and 2a). As such, dual CLEp receptor mechanisms regulate floral primordia formation in cooler environments, with CLV2/CRN-CIK signaling being necessary for primordia formation and CLV1-CIK function in primordia outgrowth being conditional due to repression by POL.

**Fig. 2:**
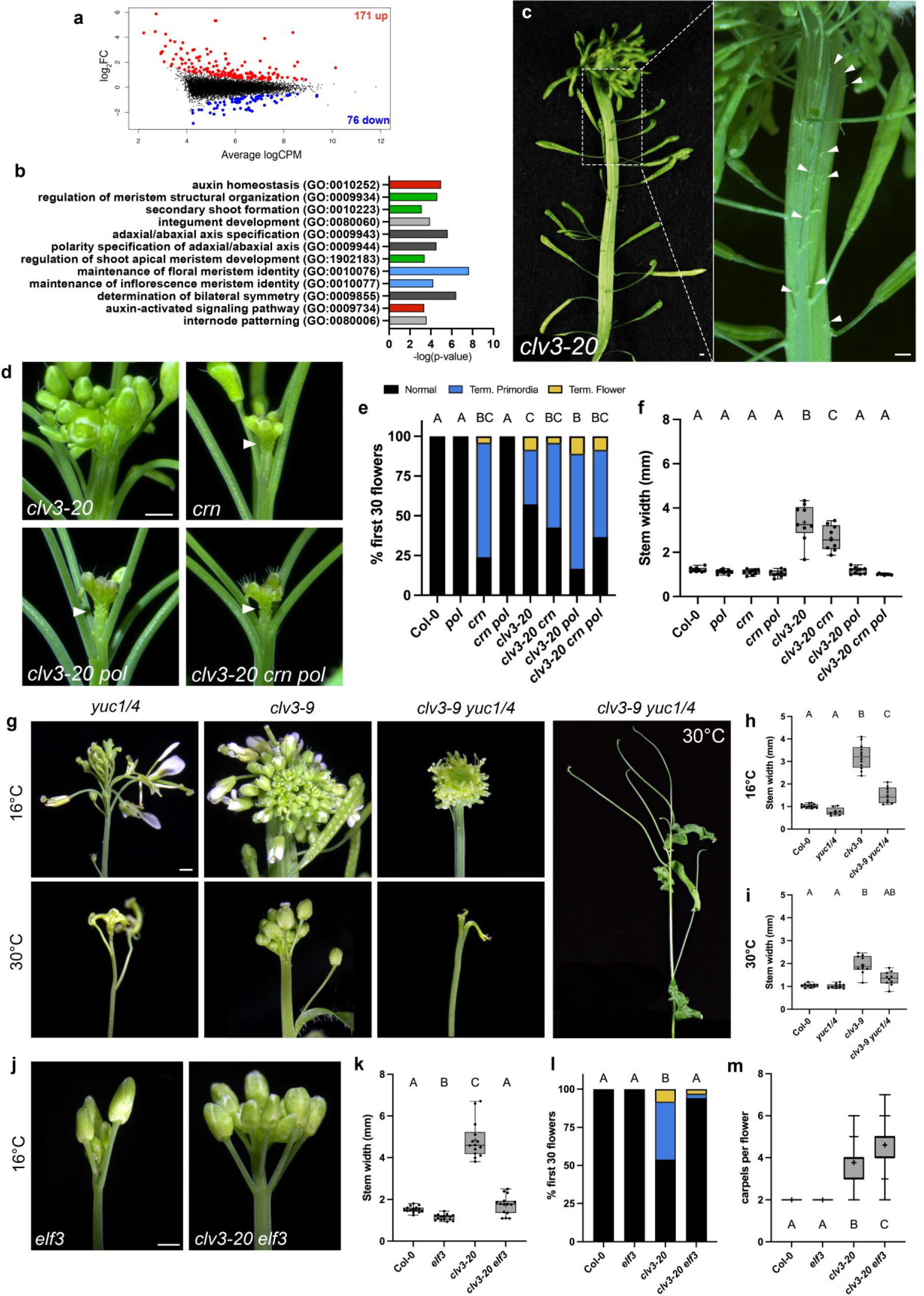
CLV3p is required for floral primordia formation across thermal environments. **a-b,** CLV3 function impacts auxin associated gene expression. **a**, Differentially expressed genes (DEGs) between *clv3-20* and Col-0 IMs with upregulated genes (red) and downregulated genes (blue) with an FDR < 5%. Black dots represent genes with FDR < 5%. **b**, GO terminology analysis showing -log(p-value) using top DEGs in *clv3-20* vs Col-0 IMs with FDR < 5% showing terms associated with auxin (red), meristem maintenance (green), flower development (blue), polarity (dark grey) and other (grey). **(c-m)** *CLV3* is necessary for floral primordia formation. **c**, *clv3-20* inflorescence side view zoomed out and close-up with arrows pointing to partially subsumed terminated flower primordia. **d**, Inflorescences of *clv3-20*, *crn*, *clv3-20 pol* and *clv3-20 crn pol* with **e**, quantification of primordia termination (n=10-22) and **f**, maximum stem widths (n=9). **g**, Inflorescence images at 16°C and 30°C of *yuc1/4*, *clv3-9* and *clv3-9 yuc14* all at same magnification. Whole plant image of *clv3-9 yuc1/4* at 30°C. Quantification of maximum stem width at 16°C (**h**, n=8-12) and 30°C (**i**, n=10). **j**, Inflorescences of *elf3* and *clv3-20 elf3* at 16°C with **k**, maximum stem width (n=14-15) and **l**, quantification of termination (n=12-15) and **m**, carpels per flower (n=10-15) at 16°C. Statistical groupings based on significant differences using Kruskal-Wallis and Dunn’s multiple comparison test correction (**e, l, m**). Maximum stem widths statistically grouped using a one-way ANOVA (**f, h, i, k**). Significance is defined as p-value < 0.05 and “+” indicates mean. Scale bars 10mm (**c**, **d, g, j**).

As de-repressed CLV1 signaling in *crn pol* plants restored primordia outgrowth we reasoned that CLV1, like CLV2/CRN, also promotes auxin outputs during primordia formation. To test this, we assayed the expression of the auxin transcriptional reporter DR5::GFP in plants grown at cooler temperatures. As expected, wild-type plants displayed strong DR5::GFP signal in incipient primordia (P0 stage^6^), which persisted through primordia outgrowth (P1), before decreasing around the P2 stage (Fig. 1d). In contrast, DR5::GFP expression in *crn* plants was almost undetectable in IMs and very low in presumptive incipient primordia sites, consistent with the lack of primordia formation (Fig. 1d). DR5::GFP expression was partially restored in *crn pol* incipient primordia, and this effect was strongly reversed in *clv1 crn pol* plants (Fig. 1d and Extended Data Fig. 1e), tracking with primordia outgrowth phenotypes (Fig. 1a,b), confirming that CLV1 promotes auxin responsiveness during primordia outgrowth.

*CLV1* is enriched in OC cells, where it is necessary and sufficient for stem cell regulation^21^ but is also expressed in L2, and occasionally L1 cells, of incipient primordia^21, 32^, while *POL* is broadly expressed in IMs (Extended Fig. 1f,g). However, in all genetic backgrounds, DR5::GFP signal was limited to L1 cells of incipient primordia. As such we asked where in the IM POL regulates CLV1- dependent primordia outgrowth by transforming *crn pol* plants with a functional *POL-YFP* transgene expressed from either the native *POL* promoter, the *WUS* promoter (OC expressed), the *CRN* promoter, or the *AINTEGUMENTA* (*ANT)* promoter (primordia expressed^46^, Fig. 1e). Consistent with previous data^21, 47^, the expression of *POL-YFP* from *POL* or *WUS* promoters, but not *CRN* or *ANT* promoters, fully restored stem cell defects in *crn pol* plants (Fig. 1f). Notably, this also mirrored the phenotypic effects on primordia formation (Fig. 1g), with *WUSpro::POL- YFP* reverting primordia outgrowth in *crn pol* plants. This suggests that POL/CLV1 signaling in OC cells promotes peripheral zone L1 cell auxin responsiveness in a non-autonomous fashion (Extended Data Fig. 1g). Interestingly, *CRNpro::POL-YFP* failed to revert *crn pol* primordia outgrowth, suggesting that CLV1 and CLV2/CRN signal from distinct IM domains to promote primordia outgrowth.

### Primordia formation and stem cell repression require CLV3p

To date no *CLE* mutant recapitulates the primordia defects found in *crn* or *clv1 crn pol* plants. CLV3p signals via both CLV1 and CLV2/CRN during stem cell regulation^14, 22^, yet flower production in *clv3* plants is notoriously robust^31, 48^. Despite this, Gene Ontology (GO) enrichment analysis of RNA sequencing data (RNAseq) from *clv3* IMs revealed subtle impacts on auxin associated genes compared to wild-type plants (Fig. 2a,b), in addition to the expected effects on stem cell and meristematic genes, suggesting CLV3p might influence auxin outputs. As such we inspected primordia formation closely in CRISPR generated *clv3-20* mutants, which eliminate CLV3p production (Extended Data Fig. 5e,f and Table 1) and found the presence of sporadic aborted primordia interspersed among the numerous normal flowers (Fig. 2c). This effect was also seen in *clv3-9* mutants, which also eliminates CLV3p production^21^ (Extended Data Fig. 5b). Aborted primordia in *clv3* mutants were often hard to resolve and appeared merged with fasciated stem tissue, suggesting fasciation in *clv3* may obscure aborted primordia. To reduce *clv3* fasciation we generated *clv3-20 pol* double mutants^44^, and found that the resulting plants now displayed numerous clearly terminated primordia comparable to *clv2*/*crn* mutants, which were not observed in *clv3* or *pol* plants (Fig. 2d-f). *clv3* also strongly reverted floral primordia formation in *crn pol* plants (Fig. 2d-f), indicating that CLV3p also promotes CLV1-dependent primordia formation. As such, *CLV3* plays a critical role in primordia outgrowth which is partially masked by stem fasciation. Heat restored primordia outgrowth in *clv3* plants, but to a lesser degree than in *crn* plants (Extended Data Fig. 3a,c). To test if heat induced primordia formation in *clv3* plants was sensitized to auxin levels, we generated *clv3 yuc1/4* triple mutants and grew these and control plants in different thermal environments. *yuc1/4* plants reduce IM auxin levels but develop interpretable shoots with identifiable flowers, unlike higher order *yuc1/2/4/6* mutants^39^. Surprisingly, *clv3 yuc1/4* plants grown in warmer conditions displayed pin-formed primary and secondary shoots which were largely devoid of flower production, something not seen in either parental genotype in warm or cooler growth conditions (Fig. 2g). As such, *CLV3* is a critical regulator of flower primordia production and synergizes with heat-induced *YUC*-mediated auxin production in warmer environments to maintain floral primordia formation.

**Fig. 3:**
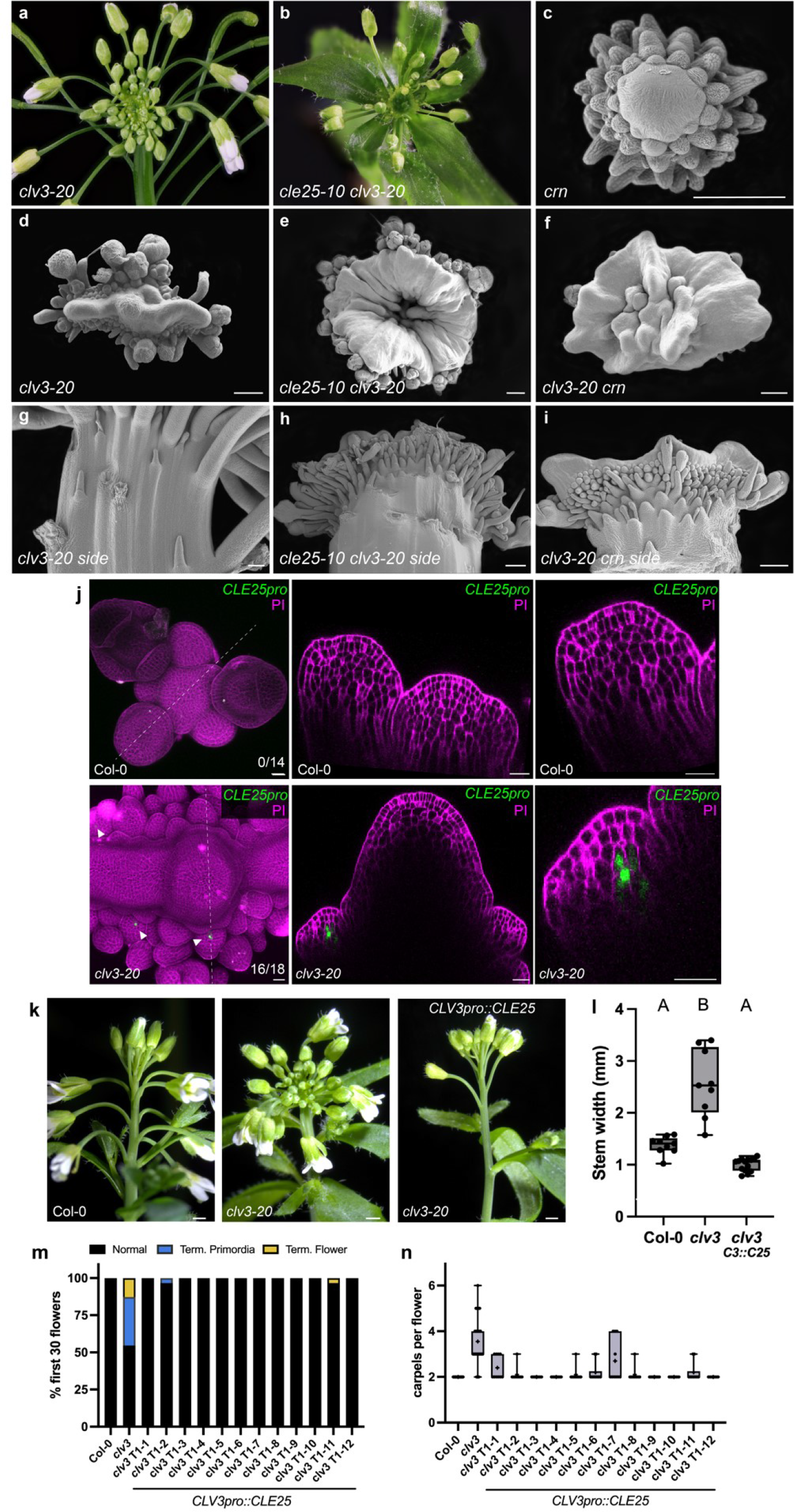
*CLV3* functions with *CLE25* in floral primordia and meristem regulation. **a-i**, *cle25 clv3-20* mutants resemble *clv3-20 crn* mutants. Images from left to right of inflorescences of *clv3-20* (**a**) and *cle25 clv3-20* (**b**). **c-f**, Aerial view SEM images of inflorescence meristems of *crn* (**c**), *clv3-20* (**d**), *cle25 clv3-20* (**e**), *clv3-20 crn* (**f**). **g-i**, Side view SEM images of inflorescence meristems of *clv3-20* (**g**), *cle25 clv3-20* (**h**), and *clv3-20 crn* (**i**). **j**, CLV3 represses *CLE25* expression in developing primordia and floral meristem. Confocal imaging of *CLE25pro::YPET-H2AX* (green) in Col-0 and *clv3-20* showing average intensity projection, axial view of IM (dotted line) and close-up axial view of developing stage 2-3 floral primordia (PI, magenta). Number of T1 lines per genotype with expression in floral primordia are indicated out of total imaged. **k**, Representative inflorescences images, **l**, maximum stem width, **m**, percent first 30 flowers and **n**, carpels per flowers of Col-0 (n=12), *clv3-20* (n=12), and *clv3-20* with the *CLV3pro::CLE25* transgene (*C3::C25*, n=12). m/n show data from individual T1 transgenics, l shows pooled data across T1 transgenic lines. Maximum stem widths statistically grouped using a one-way ANOVA (**l**). Significance is defined as p-value < 0.05 and “+” indicates mean (**l, n**). Scale bars 200µm (**c-i**), 10mm (**k**), 20µm for SEM images (**j**), with differential magnification used to highlight IM shape and details.

**Table 1:**
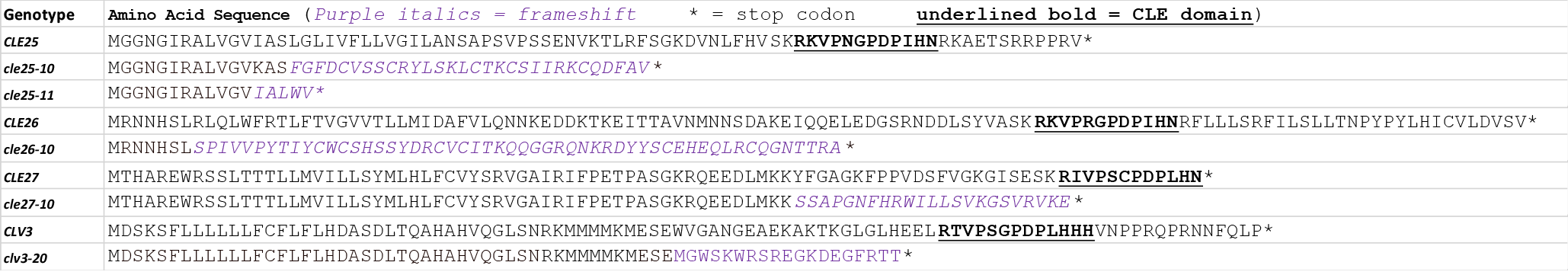
Protein sequences resulting from CRISPR CLE gene mutants used in this study. Wild-type and mutant CLE amino acid sequences are shown indicating frameshift mutations (purple italics) and early stop codons (*) before the CLE domains (underlined bold).

**Table 2:**
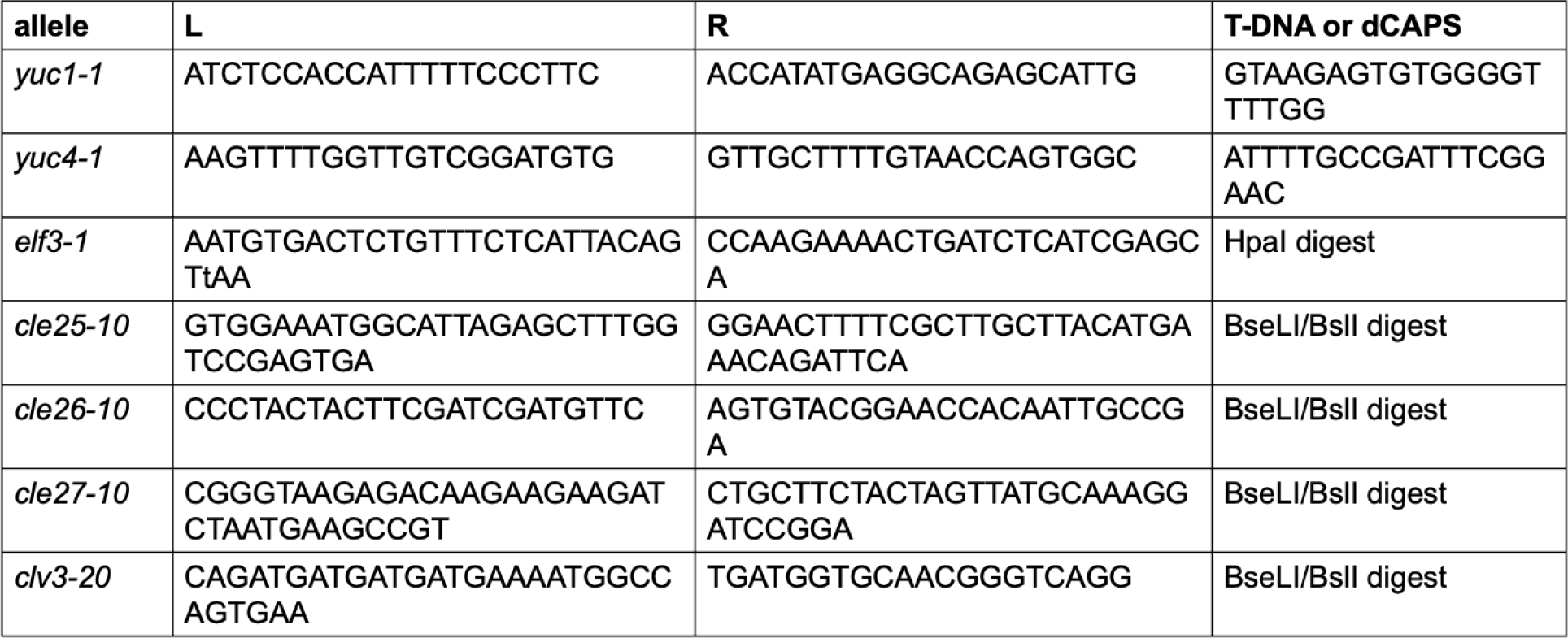
Genotyping primers generated for this study. T-DNA primers show LB, RB and T-DNA border primers. dCAPS primers shows L and R primers with enzyme used for digestion.

Given the effects on primordia formation, we next asked if heat and auxin also impacted *CLV3*- mediated stem cell regulation. Indeed, heat suppressed *clv3* stem fasciation, (Fig. 2g-i, Extended Data Fig. 3b,c) and strongly repressed carpel numbers in *clv* mutants, with stronger suppression in null *clv* signaling mutants such as *crn-10* and *clv1-101* mutants (Extended Data Fig. 4a). As such, heat and CLV signaling intersect to buffer stem cell proliferation across thermal environments. Furthermore, stem fasciation and carpel numbers were suppressed in *clv1 yuc* and *clv3 yuc* mutant combinations in ambient temperatures, demonstrating these phenotypes are also sensitized to auxin levels (Fig. 2g-i, and Extended Data Fig. 4c). Interestingly, IM expansion was enhanced in *clv3 yuc1/4* plants compared to *clv3* plants in cooler environments (Fig. 2g and Extended Data Fig. 5a), supporting previous data that IM size and auxin levels are inversely correlated ^49^, and suggesting that stem fasciation and IM expansion can be uncoupled in *clv* mutants. In cooler environments, mutations in the thermomorphogenesis repressor *elf3* suppressed *clv3* fasciation and primordia abortion (Fig. 2j-l), mirroring heat effects as expected. Surprisingly, carpel numbers were slightly increased in *elf3 clv3, elf3 clv1*, *elf3 crn*, and *elf3 clv1 crn* mutants grown in cooler temperatures compared to controls, and this trend was maintained in warmer environments despite overall lower carpel numbers (Fig. 2m and Extended Data Fig. 4b). As such, loss of *ELF3* and heat have similar buffering effects on stem fasciation and primordia outgrowth in *clv* mutants but have divergent effects on FM stem cell regulation. Collectively these data reveals that, like primordia outgrowth, CLV-dependent stem cell processes are also sensitized to auxin levels and heat, but with distinct impacts across meristem types.

**Fig. 4:**
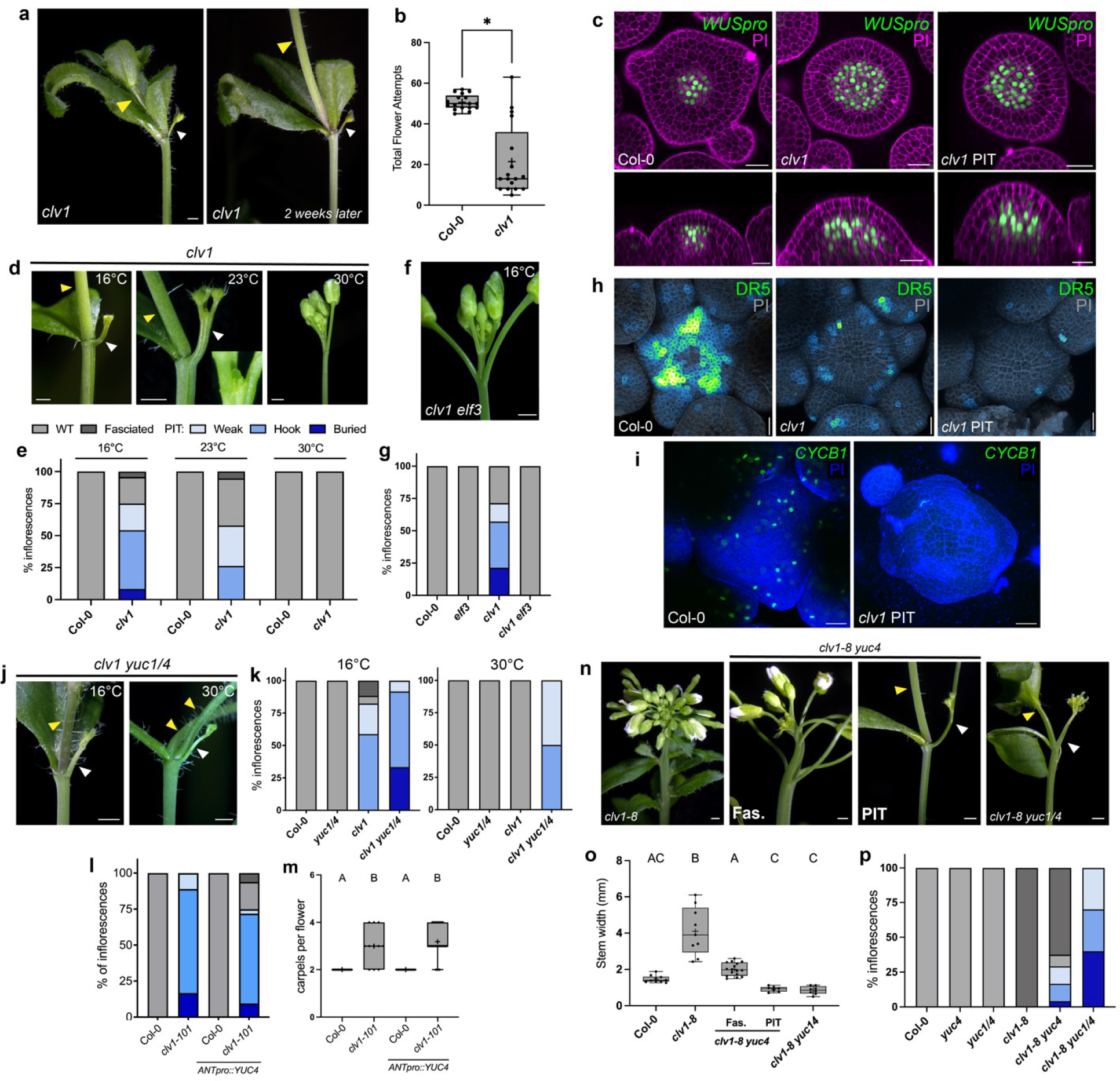
CLV1 is required for auxin-dependent IM maintenance across thermal environments. **a-c**, CLV1 signaling prevents primary inflorescence meristem termination (PIT). **a,** Inflorescences of *clv1-101* indicating primary inflorescence (white arrow) and secondary inflorescence (yellow arrow) that grows out on same inflorescence after two weeks while primary inflorescence remains terminated (PIT). **b**, Total flower attempts on primary inflorescence in Col-0 (n=17) and *clv1-101* (n=17) collected at end of life in 16°C (* indicates p-value < 0.0001). **c**, Expression of *WUSpro::YPET-N7* (green) in Col-0 (n=6), *clv1-101* (n=4) and *clv1-101* undergoing PIT (n=9) inflorescence showing L3 and axial view (PI, magenta). (**d-p**) *CLV1* promotes auxin dependent primary meristem maintenance. **d**, Inflorescences and **e**, quantification of percent inflorescences for *clv1-101* at 16°C (n=13-24), 23°C (n=16-19) and 30°C (n=12-16) that are WT-like (grey), fasciated (dark grey), weak PIT (light blue), hook PIT (blue), buried PIT (dark blue, see text). **f**, Inflorescence of *clv1-101 elf3* and **g**, PIT quantification in Col-0 (n=11), *elf3* (n=14), *clv1-101* (n=14) and *clv1-101 elf3* (n=13). **h**, Maximum intensity projections of DR5::GFP (green fire blue LUT) and PI (grey) of Col-0 (n=8), *clv1-101* (n=5) and *clv1-101* undergoing PIT (n=10). **i**, Maximum intensity projection of *CYCB1::GFP* (green) in inflorescence meristem of Col-0 and *clv1-101* PIT, PI (blue). **j**, Inflorescences of *clv1-101 yuc1/4* at 16°C and 30°C and, **k,** PIT quantification at 16°C (n=12-17) and 30°C (n=8-16) of Col-0, *yuc1/4*, *clv1-101* and *clv1-101 yuc1/4.* **l**, Quantification of percent inflorescences and **m**, carpels per flower of Col-0 (n=16), *clv1- 101* (n=18), Col-0 *ANTpro::YUC4* (n=14), and *clv1-101 ANTpro::YUC4* (n =32). Data is pooled analysis of all T1 plants from a single population experiment. **n**, Inflorescence images; **o**, maximum stem width; and **p**, PIT quantification for Col-0 (n=10), *yuc4* (n=15)*, yuc1/4* (n=10), *clv1-8* (n=10), *clv1-8 yuc4* (fasciated n=17 and PIT n=7), and *clv1-8 yuc1/4* (n=9). Temperatures as listed otherwise **a-c, f, g, i, n-p** grown at 16°C and **h** grown at 23°C. Statistical groupings based on significant differences using Kruskal-Wallis and Dunn’s multiple comparison test correction (**m**), Mann-Whitney test (**b**) and one-way ANOVA (**o**) with significance defined as p-value < 0.05 and “+” indicates mean. Scale bars 10mm (**a, d, f, j, n**), 20µm (**c, h, i**).

We suspected CLV3 was redundant with other CLE peptides, as *clv3 crn* double mutants are synergistic in cooler environments, displaying numerous terminated primordia and stalled shoots like *crn* plants, but also distinctive disk-shaped IMs, in contrast to the ribbon shape IMs typical of *clv3*^48^ for reasons unknown (Fig. 3a,d,f, and Extended Data Fig. 5d). Fortuitously, during efforts to multiplex mutate *CLE* genes using CRISPR we identified a *clv3 cle* quadruple mutant which phenocopied the distinctive IM of *clv3 crn* plants (Extended Data Fig. 5e,f and Table 1). Genetic analysis from outcrosses and independent alleles confirmed this was due to simultaneous loss of *CLE25* and *CLV3* (Fig. 3a-i and Extended Data Fig. 5a,b,c). *cle25 clv3* double mutants displayed greatly enhanced numbers of aborted floral primordia relative to *clv3* plants (Fig. 3g-i, and Extended Data Fig. 5b), as well as defective shoot elongation and disk-like IMs (Fig. 3b, d-f, and Extended Data Fig. 5a,c,d). Despite this, *cle25* single mutants were wild type in appearance and did not enhance *crn* like *clv3* (Extended Data Fig. 5h,i). Additionally, *CLE25* transcriptional reporter expression was not seen in wild type IMs or incipient floral primordia (Fig. 3j and Extended Data Fig 5k) but was observed in seedling root phloem as previously reported^50^ (Extended Data Fig 5j). In contrast, *CLE25* displayed sporadic ectopic expression in L3-L4 cells in stage 2-3 floral primordia in *clv3* plants (Fig. 3j and Extended Data Fig 5k), suggesting CLV3p represses *CLE25* expression. Notably, expression of *CLE25* from the native *CLV3* promoter was sufficient to restore floral primordia formation in *clv3* mutants (Fig. 3k,m) and suppress stem width and carpel numbers in *clv3* mutants. (Fig. 3l,n). As such, CLV3p promotes auxin mediated floral primordia formation and IM homeostasis in conjunction with *CLE25* as part of a transcriptional compensation mechanism akin to *CLV3/CLE9* compensation in tomato^23^.

### Auxin-dependent meristem maintenance in cool growth environments requires CLV1

Our data reveals that CLV3p regulates multiple temperature-sensitive IM functions simultaneously through partially overlapping receptors, complicating the analysis of individual IM processes in *clv3* mutants. As such, we wondered if other CLEp receptors had subfunctionalized IM roles which are occluded in *clv3* plants and warmer environments. Indeed, careful inspection of *clv1-101* null mutants revealed a novel phenotype unique to *clv1* that we termed Primary Inflorescence Termination (PIT) which was observed in ambient temperatures but was more penetrant and severe in cool environments. During PIT, the *clv1* primary shoot generates a few normal flowers and then all primordia formation and outgrowth ceases and primary shoot elongation aborts (Fig. 4a,b and Extended Data Fig. 6a,b). Unlike *crn* plants, which form terminated floral primordia and recover, *clv1* plants that undergo PIT cease all apparent IM activity and never recover (Fig. 4a). As PIT penetrance and severity varied, we classified PIT phenotypes by decreasing severity into “buried” (primary shoot undergoes PIT before emerging fully from rosette); “hook” (primary shoot elongates but undergoes PIT upon secondary branch formation resulting in a “hook”-like primary shoot); and “weak” classes (like hook class, but with additional primary shoot elongation and flower formation before PIT, Extended Data Fig. 6b). Quantification of *clv1* null mutants revealed that over 80% of plants underwent PIT and failed to form a normal shoot in cool temperatures (Extended Data Fig. 6c), and similar PIT was observed in the *clv1-20* knock-down allele^51^ (Extended Data Fig. 6e,f). Despite this, *clv1* PIT IMs retained the dome shape of *clv1* mutant IMs, contained apparently normal IM cell layer organization (L1, L2, OC), and expressed *WUS* appropriately in OC cells (Fig. 4c), but displayed little expression of the *CYCLINB1;2* G2/M cell cycle marker^52^(Fig. 4i). As such PIT is not associated with a loss of *WUS*- expression or gross defects in IM structure but instead appears correlated with a global cessation of mitotic activity across all IM zones. Strikingly, PIT phenotypes were not observed in warm temperatures (Fig. 4d,e), or in *elf3 clv1* double mutants grown in cool/ambient temperatures (Fig. 4f,g), reminiscent of heat and *elf3* effects on primordia formation in *crn* plants. This suggested that PIT might also be linked to lowered auxin functions in cooler temperatures. Consistent with this *clv1* IMs undergoing PIT in cool environments displayed nearly undetectable levels of DR5::GFP signal (Fig. 4h) and elevated levels of the DII-Venus auxin perception reporter^53^ (Extended Data Fig. 6d). Additionally, partial reduction in IM auxin by combining *yuc1/yuc4* mutations with *clv1- 101* enhanced PIT penetrance and severity in cooler environments, and conversely *YUC1/4* were necessary for full heat mediated PIT suppression, strongly suggesting that low IM auxin levels are causal to *clv1* PIT (Fig. 4j,k). Indeed, expression of *YUC4* from the *ANT* promoter repressed the severity and occurrence of PIT in *clv1* plants (Fig. 4l), without altering carpel number (Fig. 4m). Occasionally we also observed fasciation in *clv1-101* null mutants and interestingly these plants very rarely underwent PIT. This phenotypic variability suggests that null *clv1* mutants are poised at a threshold of stem cell regulation in which subtle perturbations can give rise to different developmental states. Correspondingly, strong *clv1-8* alleles always fasciated and never displayed PIT (Fig. 4n-p), and PIT was also not observed in fasciated *crn clv1-101* or *clv3* plants, suggesting fasciation masks PIT as it does *clv3* primordia defects. Supporting this, *clv1-8 yuc4* double mutants and *clv1-8 yuc1/4* triple mutants displayed significantly reduced fasciation as expected (Fig. 4n,o), and this correlated with the appearance of PIT in *clv1-8 yuc1/4* triple mutants and also in *clv1-8 yuc4* double mutants, where it was mutually exclusive with fasciation (Fig. 4n-p). Collectively these results indicate that CLV1 plays a novel and critical role in auxin dependent IM maintenance across thermal environments which is partially masked by IM fasciation and heat induced auxin biosynthesis.

## Discussion

Understanding how plants integrate diverse cues to maintain development across environments is a key challenge. Our work reveals that temperature and CLV signaling intersect to maintain multiple auxin-regulated IM functions across thermal environments (Fig. 4n), including classical CLV-stem cell processes. CLV3p plays an unappreciated and critical role in auxin-dependent floral primordia generation and CLV1 and CLV2/CRN receptors can both act in this process, explaining previous enigmatic observations on flower formation in *CLV*-class mutants^21, 54–56^. As such, CLV3/CLE signaling can repress and promote cell proliferation in different IM domains through the same suite of receptors. Whether this reflects conserved or divergent downstream signaling outputs is unclear. CLV1 appears to promote L1 auxin responsiveness cell non-autonomously from OC cells, but not *CRN* expressing cells, suggesting spatial differences in CLEp receptor IM expression could be important. However, CLV3p diffuses broadly in meristems^24^, and is expressed in the central zone 3-5 cells away from P0/P1 stage primordia^57^, suggesting it could influence primordia directly. *CLV3* function in primordia generation is partially buffered by transcriptional compensation from *CLE25* in emerging primordia, with CLV3 repressing *CLE25* expression in primordia and ectopic *CLE25* thereby partially compensating for loss of *CLV3*. As such, Arabidopsis IM functions are buffered by *CLV3*/*CLE* transcriptional compensation as in tomato^23^, in addition to *CLV1/BAM* receptor transcriptional compensation^21, 22^. *CLE25* buffering impacts floral primordia formation and IM expansion but does not appear to influence floral stem cell regulation as *cle25* mutants do not enhance *clv3* carpel numbers (Extended Data Fig. 5g). This likely reflects temporal or spatial differences in *CLE25* expression as *CLE25* expressed from the *CLV3* promoter can restore *clv3* carpel numbers.

How CLEp signaling modulates auxin outputs is unknown. The reversion of *clv1* PIT and primordia defects by heat or loss of *ELF3*, and the rescue of primordia defects in *crn* by ectopic *YUC1*^31^, suggests that CLV signaling is not critical for auxin perception, transport, or signaling. *WUS* has been proposed to act as an auxin “rheostat”^58^, however it is not clear if WUS mediates *clv1/crn* auxin processes as *crn* primordia defects are not associated with ectopic *WUS* expression and there are no differences in *WUS* expression in *clv1* plants undergoing PIT versus those that do not. Furthermore, ectopic *WUS* misexpression does not recapitulate PIT or primordia phenotypes^17, 21^, and extreme fasciation masks rather than enhances PIT and primordia phenotypes. Interestingly, moss *clv* mutants also display altered auxin sensitivity^59^, suggesting CLV-auxin interactions may have an ancient origin which could pre-date *WUS* evolution^60^. While IM size and auxin levels appear inversely correlated, supporting previous models^49^, the PIT phenotype suggests CLV1 has a novel role in promoting auxin-dependent IM maintenance. The precise nature of PIT is unclear, but it is associated with a global cessation of IM functions in the absence of gross IM patterning change, perhaps explaining why it is obscured by IM over-proliferation in some backgrounds. PIT phenotypes overlap with *crn* shoot and primordia defects to a degree, but key differences exist suggesting they are not equivalent. Despite this, PIT is sensitized to temperature and auxin levels, like other CLV IM processes, reminiscent of heat and *YUC* impacts on *de novo* shoot regeneration^61^. Interestingly, our data suggests heat and loss of *ELF3* do not appear entirely equivalent. Both conditions repress PIT, floral primordia termination, and IM fasciation phenotypes. However, heat represses *clv* carpel numbers and loss of *ELF3* slightly enhances them, suggesting differential wiring in FM and IM processes. Curiously, heat enhances the primordia outgrowth defects in *clv3 yuc1/4* mutants, similar to but more severe than *clv2 yuc1/yuc4* primordia defects^31^. This is reminiscent of the increased penetrance of auxin mediated seedling defects seen in warmer temperatures^62^, suggesting that heat itself may negatively impact auxin function. Understanding how CLEp signaling, auxin, and temperature impact the diverse functions of shoot meristems could inform the creation of climate change resistant crops.

## Methods

### Plant materials and growth conditions

All Arabidopsis (*Arabidopsis thaliana*) mutant and transgenic lines used were in the Columbia-0 (Col) ecotype. The following lines were used and described previously: *crn-10*^22^*, pol-6*^43^*, clv1- 101*^63^*, clv1-8*^64^*, clv1-20*^51^*, cik1-1*^36^*, cik1-3*^31^*, cik2-1*^36^*, cik2-3*^31^*, cik3-1*^36^*, cik4-1*^36^*, cik4-3*^31^*, clv3- 9*^21^*, yuc1-1*^39^ (SALK_106293)*, yuc4-1*^39^ (SM_3_16128)*, elf3-1*^65^(ABRC CS3787).

*CYCLINB1;2pro:Dbox-GFP* in the Col-0 background was a gift from Cristina Ferrándiz^52^ and was introgressed into the *clv1-101* background. Seeds were sterilized and plated on 0.5% Murashige and Skoog (MS) medium buffered with MES (pH 5.7). Plates were stratified for 2 days at 4°C and moved to a constant light growth chamber at 22°C. After 7 days seedlings were transplanted to soil and moved to one of three temperatures with continuous light; 16-17°C growth chamber (cool temperature), 22-23°C grow room (ambient temperature) or 30°C growth chamber (warm temperature). Plants are grown in ambient temperature unless otherwise stated. *WUSpro::YPET- N7* in Col-0^31^, *DR5pro::GFP* in Col-0, and *CLV1pro::2xYPET-N7* in Col-0^21^ lines were crossed into additional alleles for analysis.

### Confocal microscopy and image analysis

Live *Arabidopsis* inflorescence meristem imaging was performed as previously reported using an inverted Zeiss 710 confocal laser scanning microscope with an inverter attachment^66^. Floral transitioned IMs were dissected carefully, and briefly stained with PI (final concentration between 50μg/mL for WT and 150μg/mL for *clv* mutants) on ice. 2-3mm of the IM stem was vertically inserted in 1.5-2% agarose (w/v) in a Petri dish and immersed in cold water, aspirating away air bubbles formed around the IM with a pipette. IMs were imaged using the W Plan-APOCHROMAT 40X (NA = 1.0) water dipping objective. Laser excitation and detected emission are as follows: Ypet markers with PI - 514nm argon laser, Ypet channel 520-581nm, PI channel 655-758nm; GFP markers with PI - 488nm argon laser, GFP channel 493-556nm, PI channel 598-642nm argon laser. Images comparing reporter levels in different backgrounds were imaged with identical specifications in the ZEN software and post-processed with identical specifications using Fiji/ImageJ (version 1.0, NIH) to generate single scan images and axial views of IMs. *DR5::GFP* and *CIKpro:YPET-H2AX* were represented with green fire blue LUT (Fiji).

### Scanning Electron Microscopy

Scanning electron microscopy of unfixed, hydrated tissue was performed using a Hitachi TM4000Plus II SEM with motorized stage set to Charge Reduction vacuum mode. Young inflorescences were dissected to remove open flowers and older buds and to expose the IM. Dissected samples were immediately mounted with double-sided conductive carbon adhesive (25mm OD PELCO Tabs™ Ted Pella, Inc.) onto the sample holder. Samples were processed quickly from dissection to image acquisition to minimize shrinkage. Images were edited using Adobe Photoshop (v23.3.1) to remove the background.

### Photography

Unless otherwise specified, images of terminating inflorescences and controls were acquired using a Zeiss Stemi 2000C microscope equipped with a Zeiss Axiocam 105-color digital camera and Zeiss Zen software. Composite images of inflorescences from *clv3 yuc1/4* (Fig. 2g, far right panel) *clv3* (Fig. 3a), and *cle25 clv3* (Fig. 3b) were made using Affinity Photo (version 1.6.5.135) Focus Merge tool and edited for contrast. Images were photographed using a Canon Eos Rebel T6i DSLR camera on a tripod equipped with a Tamron 60mm f/2.0 SP DI II LD IF Macro lens (Fig. 2g), or a Canon EF-S 35mm f/2.8 Macro IS STM lens (Fig. 3a,b).

### Plasmid Construction and Generation of Transgenic Lines

All binary vectors were introduced into *Agrobacterium tumefaciens* strain GV3101 by electroporation and transformed into plants using the floral dip method^67^. The *POLpro::POL-YPET, CRNpro::POL-YPET, ANTpro::POL-YPET* and *WUSpro::POL-YPET* were generated using Gateway technology (Thermo) with existing entry *(pENTR-D POL-YPET)* and binary vectors in the pMOA33 and pMOA34 background^68^ *(MOA34POLpro::GTW, MOA33CRNpro::GTW, MOA34ANTpro::GTW, MOA34WUSpro::GTW)* that were transformed into *pol crn* backgrounds and plated on either Kanamycin *(MOA33)* or Hygromycin *(MOA34)* 0.5% MS plates. Transcriptional reporter for *POL* was generated using *POLpro::GTW* and an entry vector generated using TOPO cloning, *pENTR::YPET-H2AX* through gateway technology. Transcriptional reporters for *CIK1/2/3/4* were generated by either PCR amplifying 3kb upstream of *CIK1* and 2.6kb upstream of *CIK2* and TOPO cloning into *pENTR-D* (ThermoFisher) or synthesis of 3kb upstream of *CIK3* and 1.7kb upstream of *CIK4* cloned into a gateway compatible entry vector (Twist biosciences). These entry vectors were gateway cloned into the *YPET-H2AX GTW* (Gift from Kevin Potter) and transformed into Col-0 plants. The transgene was selected on Kanamycin plates and 4-6 lines were screened for consistent expression patterns. *CLE25pro::YPET-H2AX* was generated by the synthesis of 2.5kB upstream of *CLE25* into a gateway compatible entry vector (Twist bioscience) that was cloned into the *YPET-H2AX GTW* vector and transformed into Col-0 and *clv3-20* plants. T1s were selected off Kanamycin plates and 6 lines were screened per genotype for expression. T1 line #8 in the *clv3-20* background was backcrossed to Col-0 wildtype for imaging of same line in different backgrounds (Extended Data Fig. 5f). *CLV3pro::CLE25* lines were generated by amplifying the *CLE25* CDS from Col-0 RNA, TOPO cloned into *pENTR-D* (ThermoFisher) and gateway cloned into a *MOA33CLV3pro::GTW* vector. This vector was then transformed into Col-0 and *clv3-20* plants and T1 plants were selected on Kanamycin plates. The *YUC4* CDS was amplified from Col-0 RNA and TOPO cloned into *pENTR-D* (ThermoFisher). This was then recombined in *MOA34ANTpro::GTW* and transformed into *clv1-101* plants. T1 plants were selected on hygromycin. *MOA33UBQ10::DII-Venus* vector was dipped into Col-0 and *clv1-101* plants and screened on Kanamycin with T1 images imaged. One T1 *clv1-101* single insert *MOA33UBQ10::DII-Venus* line was crossed to Col-0 and imaged in the F1.

### RNA Sequencing and Data Analysis

45-50 inflorescence meristems for each of three biological replicates of Col-0 and *clv3-9* were collected and total RNA isolated using the EZNA Plant RNA kit (Omega Bio-tek). RNA was treated with RNase-free DNase (Omega Bio-tek) and 1.5 ug RNA was used for library preparation at the High-throughput Sequencing Facility at UNC Chapel Hill using the Stranded mRNA-Seq kit (Kapa Biosystems. The NovaSeq 6000 sequencer (Illumina) generated 50bp paired-end reads with a read depth of 23-35 million reads per biological replicate. Pre-trimmed raw data was aligned to the *A. thaliana* reference genome (TAIR10.1) using HISAT2 version 2.2.0^69^. Reads were counted using Subread version 1.5.1^70^ and transferred to RStudio (v.1.4.1106) to be normalized using EDASeq version 2.22.0^71^ and RUVseq version 1.22.0^72^ (upper quartile normalization). Differentially expressed genes (DEGs) were identified with FDR <5% using EdgeR version 3.33.0^73^ and these top DEGs were used for GO term analysis from Panther^74^. GO terms with highest fold enrichment categories were graphed based on -log(p-value).

### CRISPR mutagenesis of *CLV3* and *CLE25*/*26*/*27*

Generation of multiplex mutants of *CLE* genes using the *pCUT* vector CRISPR system was used to create the *clv3-20*, *cle25-10*, *cle25-11, cle26-10*, *cle27-10* mutations in Col-0 plants. *U6pro::gRNA* constructs targeting *CLE25*, *CLE26* or *CLE27* genes were synthesized as a single unit by GeneArts (Regensburg, Germany) and cloned into the *pCUT4* binary vector as described in^75^. Hygromycin resistant plants were selected in the T1 generation and target genes were sequenced by Sanger sequencing and screened for mutations in the T2 and T3 generations as needed. Quadruple fixed mutants were isolated and the Cas9 *pCUT* transgene was segregated out from these plants before genetic analysis.

### Quantification and Statistical Analysis

All quantitative data was compiled and analyzed on GraphPad Prism v.9.3.1. Two to five biological replicates were performed for each experiment ensuring consistent results and representative replicate data is shown. Sample sizes are indicated in figure legends. Flower termination in first 30 flowers across genotypes and/or transgenic lines was compared using only % flower primordia termination and was compared across samples using a non-parametric Kruskal-Wallis and a Dunn’s multiple comparison test correction where significance was defined as p-value < 0.05. Carpel number across genotypes was obtained by counting 10 consecutive flowers on the primary inflorescence. For mutant inflorescences with termination, carpels were counted after recovery of flower outgrowth and for inflorescences without termination, counting started at the 6^th^ carpel. Carpel number was compared statistically using a non-parametric Kruskal-Wallis and a Dunn’s multiple comparison test correction where significance was defined as p-value < 0.05. Stem width was measured using digital calipers (SPI 6”) taking multiple measurements along primary inflorescence to identify maximum stem width. Maximum stem width was compared statistically using a one-way ANOVA if normally distributed or using a non-parametric Kruskal-Wallis and a Dunn’s multiple comparison test correction without normal distribution where significance was defined as p-value < 0.05. Comparison of Col-0 and *clv1-101* total flower attempts were done with the Mann-Whitney test where significance was defined as p-value < 0.05. Statistical comparisons of carpel number of genotypes between temperatures were done using the Mann-Whitney test where significance was defined as p-value < 0.05.

## Acknowledgments

The authors thank Tony D. Perdue, director of the University of North Carolina-Chapel Hill Genome Sciences Microscopy Core, for assistance with confocal imaging and Andrew Willoughby for assistance with SEM and photography. We thank Jamie Winshell and James Garzoni for lab and plant growth facility support. We thank UNC’s High-Throughput Sequencing Facility for sequencing services. We thank members of the Nimchuk lab for critical feedback on this project. This research was supported by a National Institute of General Medical Sciences– Maximizing Investigators’ Research Award from the NIH (R35GM119614 to Z.L.N) and National Science Foundation (NSF) Plant Genome Research Program (PGRP) (IOS-1546837 to Z.L.N). D.S.J. was supported by NSF Postdoctoral Research Fellowship in Biology through the PGRP (NSF 1906389).

## Author contributions

Conceptualization: A.J., E.S.S., D.S.J., Z.L.N; Methodology: A.J., E.S.S., D.S.J., C.L.S, Z.L.N;

Validation: A.J., E.S.S., D.S.J., C.L.S; Formal analysis: A.J., E.S.S., D.S.J., C.L.S; Investigation:

A.J., E.S.S., D.S.J., C.L.S; Data curation: A.J., E.S.S., D.S.J.; Writing (original draft): A.J., E.S.S.,

D.S.J., Z.L.N.; Writing (review and editing): A.J., E.S.S., D.S.J., Z.L.N.; Visualization: A.J., E.S.S., D.S.J.; Supervision: Z.L.N.; Project administration: Z.L.N.; Funding acquisition: Z.L.N.

The authors declare no competing interests.

Correspondence and requests for materials should be addressed to

---

**Extended Data Fig. 1:**
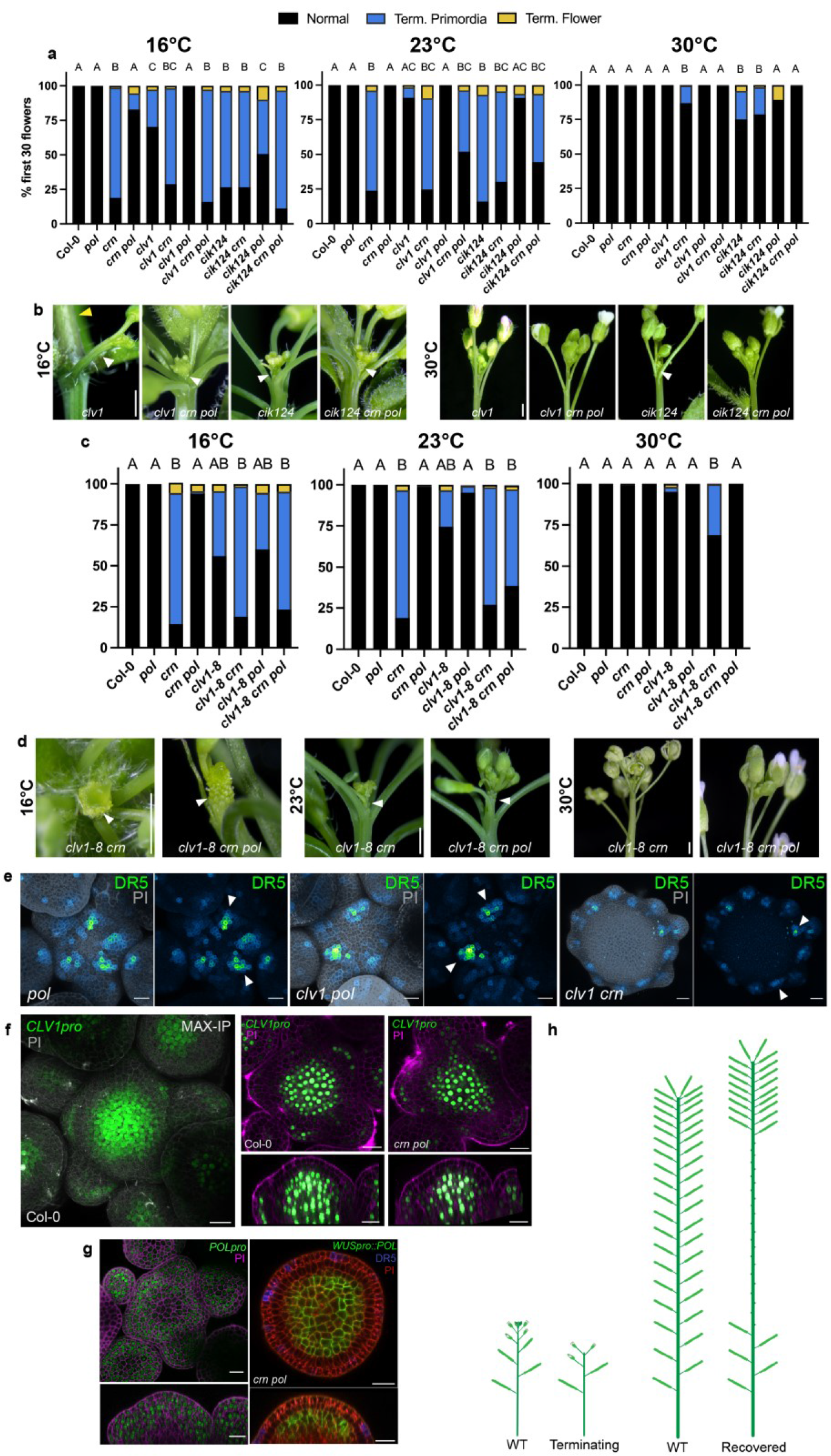
Temperature and CLV3p receptors intersect to buffer auxin mediated floral primordia formation. **a-e**, CLE receptors and temperature intersect to buffer floral primordia formation. **a,** Quantification of flower primordia termination at 16°C (n=9-21), 23°C (n=10-24) and 30°C (n=12- 15) for Col-0, *pol, crn, crn pol, clv1-101*, *clv1-101 crn, clv1-101 pol, clv1-101 crn pol*, *cik124, cik124 crn, cik124 pol, cik124 crn pol.* **b**, Inflorescence images at 16°C and 30°C of *clv1-101*, *clv1-101 crn pol*, *cik124*, and *cik124 crn pol*. All *cik* alleles in this figure are *cik1-3, cik2-3, and cik4-3* CRISPR generated alleles. Same magnification within temperatures. **c**, Quantification of flower primordia termination at 16°C (n=8-9), 23°C (n=8-13), and 30°C (n=12) for Col-0, *pol*, *crn*, *crn pol*, *clv1-8 crn*, *clv1-8 pol*, *clv1-8 crn pol*. **d**, Inflorescence images at 16°C, 23°C and 30°C of *clv1-8 crn* and *clv1-8 crn pol.* Same magnification within temperatures. **e**, Maximum intensity projection of *DR5::GFP* (green fire blue LUT) and PI (grey) and *DR5::GFP* alone in the IMs *pol* (n=7), *clv1-101 pol* (n=6), and *clv1-101 crn* (n=8). **f**, Maximum intensity projection of Col-0 *CLV1pro::2xYPET-N7* (green) inflorescence meristem. Expression patterns of *CLV1pro::2xYPET-N7* in Col-0 (n=4) and *crn pol* (n=5) showing no expression changes. **g**, Expression patterns of *POLpro::YPET-H2AX* (green, n=6) in Col-0 and *WUSpro::POL-YPET* (green, n=3) and *DR5::GFP* (purple) in *crn pol* showing L5 and Z axis (PI, red). **h**, Diagram of flower/fruit production on primary inflorescence in wild-type and terminating and recovering *crn* plants. Statistical groupings based on significant differences using Kruskal-Wallis and Dunn’s multiple comparison test correction (**a, c**) where significance is defined as p-value < 0.05. Scale bars 10mm (**b, d**) and 20µm (**e, f, g**).

**Extended Data Fig. 2:**
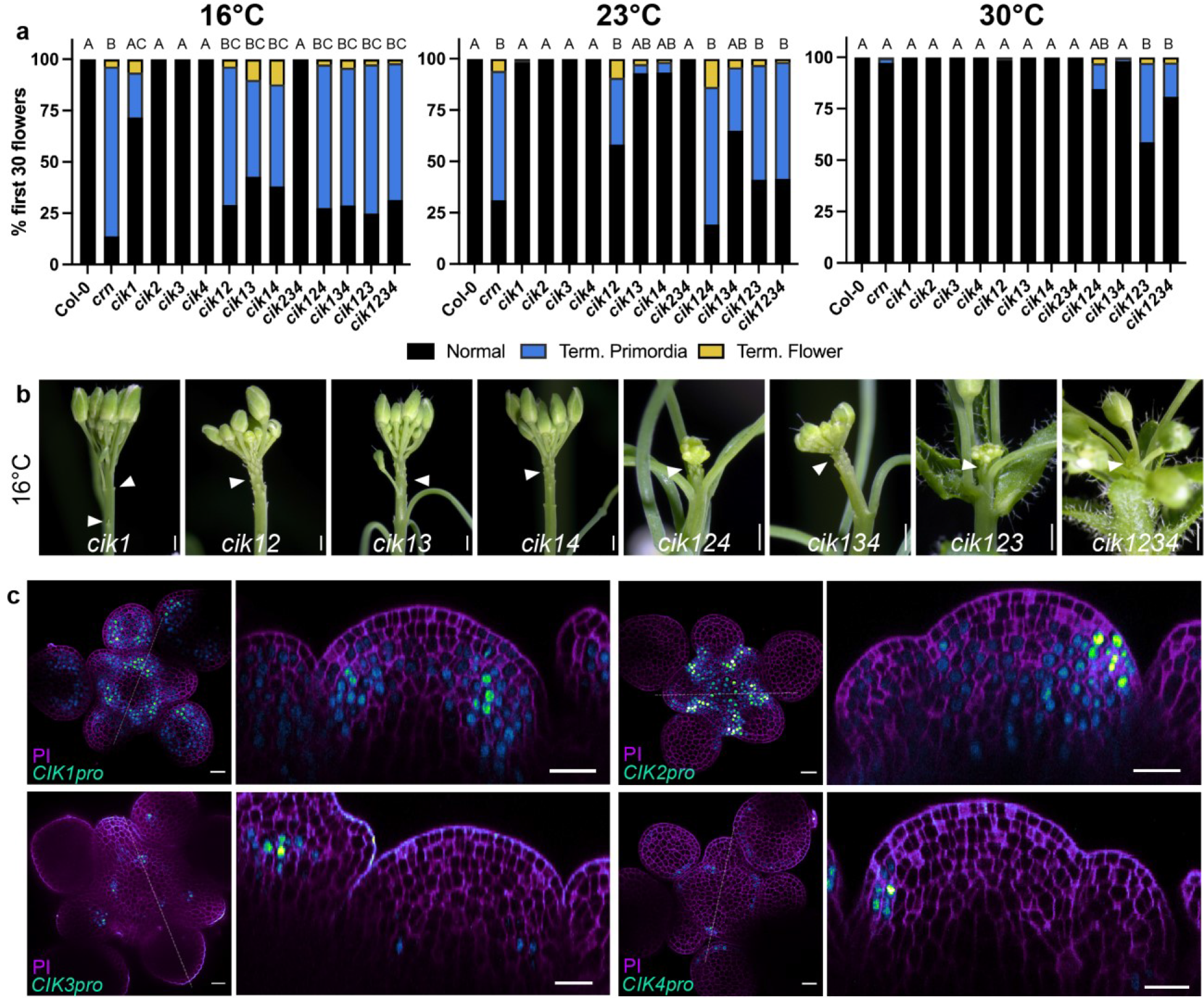
CIK co-receptors are required for temperature-dependent floral primordia outgrowth. **a-b**, CIK receptors act redundantly in temperature sensitive floral primordia formation. **a**, Quantification of flower primordia termination at 16°C (n=7-14), 23°C (n=8-9) and 30°C (n=7-9) for Col-0, *crn, cik1*, *cik2, cik3, cik4,* cik12, *cik13*, *cik14*, *cik234*, *cik124*, *cik134*, *cik123* and *cik1234*. **b**, Inflorescence images at 16°C of *cik1*, *cik12*, *cik13*, *cik14*, *cik124*, *cik134*, *cik123*, *cik1234* with arrows pointing to termination. Same magnification within single and double mutants as well as within triple and quadruple mutants. All *cik* alleles in this figure are *cik1-1, cik2-1, cik3- 1, and cik4-1* T-DNA lines. **c**. *CIK* genes are expressed in the IM. Expression patterns of *YPET- H2AX* (green fire blue LUT) reporter lines in the IM with XY view of L5 and axial view of same stack indicated with dotted line of *CIK1pro* (n=10), *CIK2pro* (n=15), *CIK3pro* (n=8) and *CIK4pro* (n=14) in Col-0 showing representative images (PI, magenta). Statistical groupings based on significant differences using Kruskal-Wallis and Dunn’s multiple comparison test correction (**a**) where significance is defined as p-value < 0.05. Scale bars 10mm (**b**) and 20µm (**c**).

**Extended Data Fig. 3:**
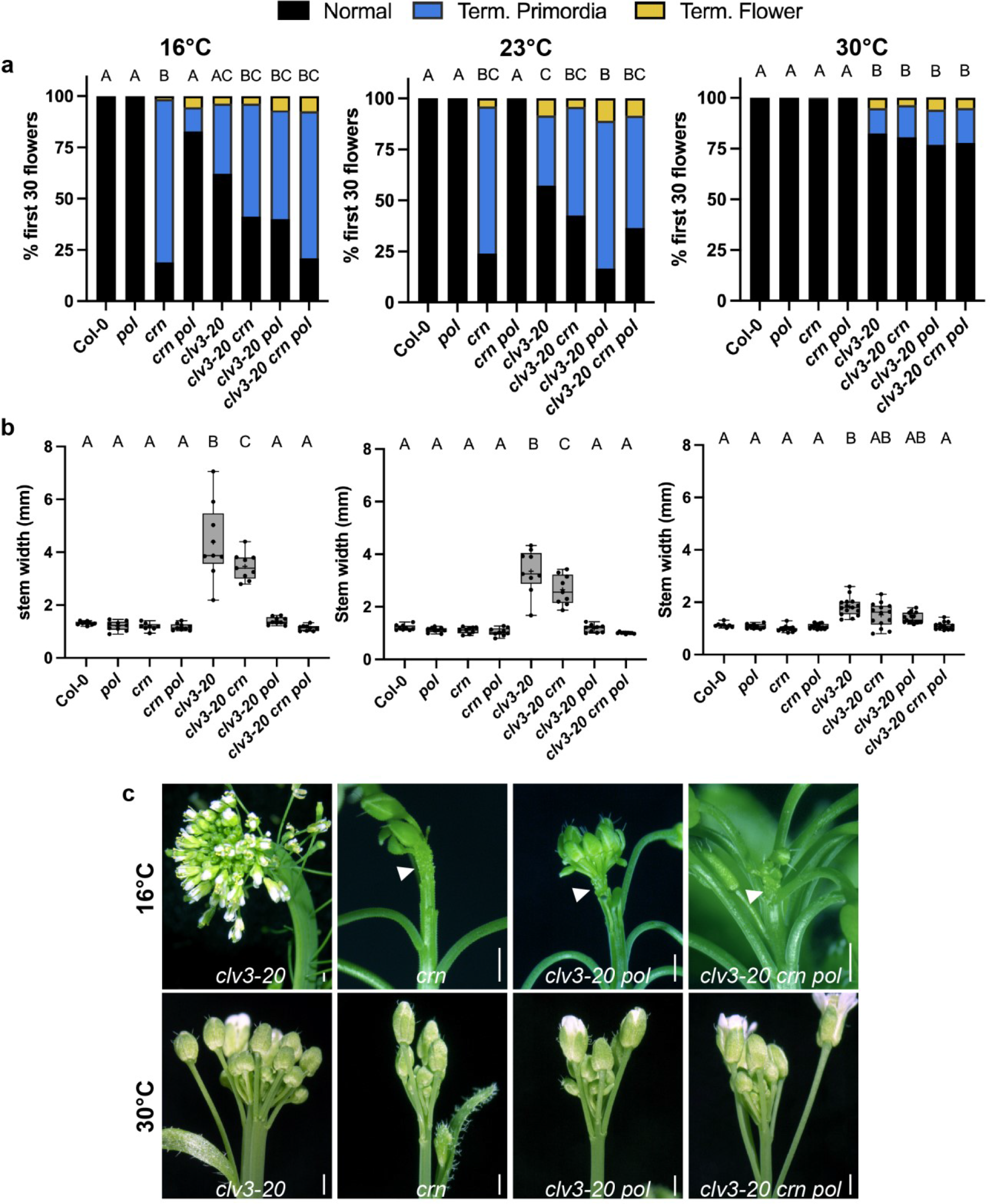
CLV3p dependent stem fasciation and floral primordia outgrowth are temperature dependent. **a**, Primordia outgrowth defects in *clv3* plants are buffered by temperature. Quantification of termination at 16°C (n=8-21), 23°C (n=9-22), and 30°C (n=12-16) for Col-0, *pol*, *crn*, *crn pol*, *clv3-20*, *clv3-20 crn*, *clv3-20 pol*, *clv3-20 crn pol*. **b**, Stem fasciation in *clv3* plants is buffered by temperature. Quantification of maximum stem width at 16°C (n=9), 23°C (n=9), and 30°C (n=15). **c**, Inflorescences at 16°C and 30°C of *clv3-20*, *crn*, *clv3-20 pol* and *clv3-20 crn pol*. Varying magnification is used at 16°C to show *clv3* fasciation and flower termination (arrows). All same magnification at 30°C. Statistical groupings based on significant differences using a one-way ANOVA (**a**, 16°C and 23°C) and Kruskal-Wallis and Dunn’s multiple comparison test correction (**a** 30°C and **b**) where significance is defined as p-value < 0.05 and “+” indicates mean. Scale bars 10mm (**c**).

**Extended Data Fig. 4:**
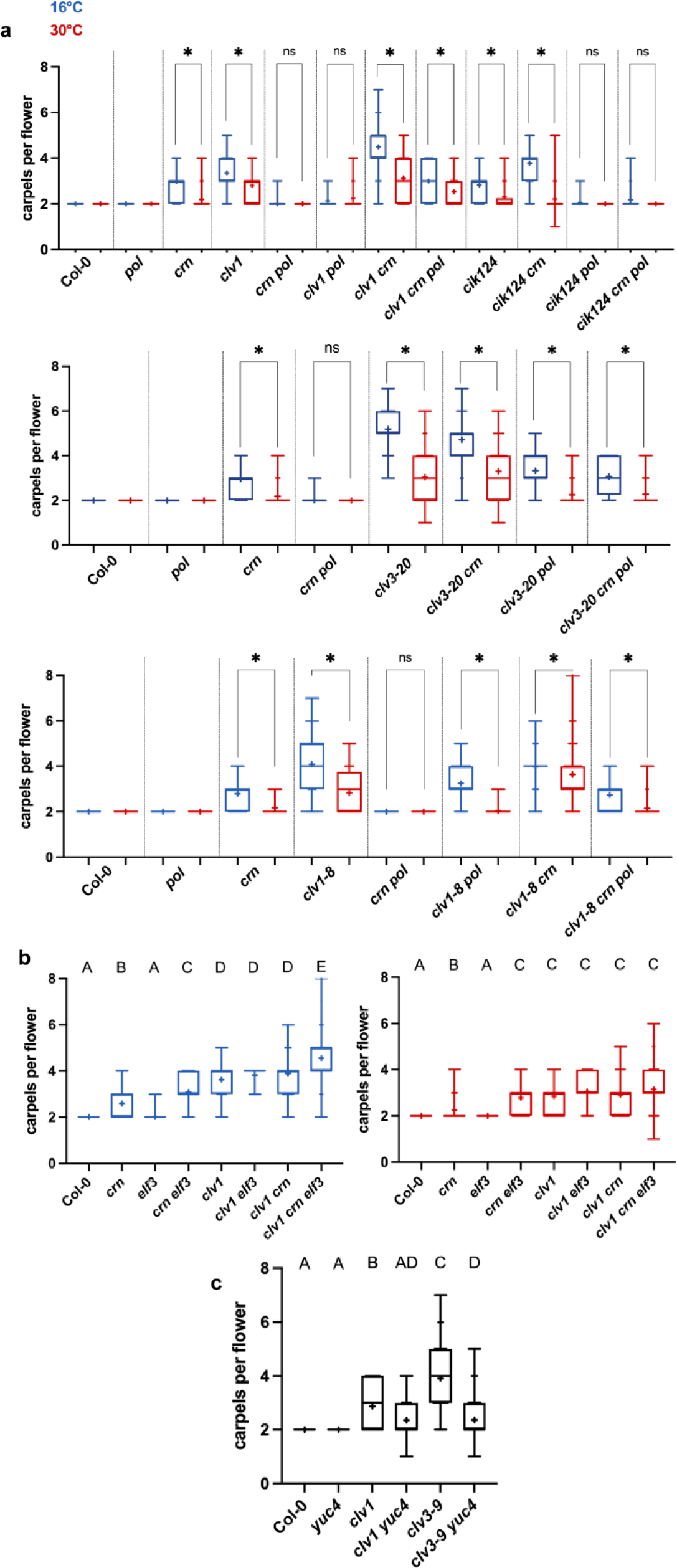
CLV-dependent FM stem cell regulation is sensitized to heat and auxin levels. **a**, *CLV*-dependent FM stem cell regulation is repressed in warmer growth environments. Carpels per flower at 16°C (blue) and 30°C (red) for *clv1-101 crn pol*, *cik1-3 2-3 4-3 crn pol*, *clv3-20 crn pol*, *clv1-8 crn pol* and all single mutant control alleles. **b**, *elf3* enhances carpel numbers in *clv1- 101* mutants. Carpels per flower at 16°C (blue) and 30°C (red) for *clv1-101 elf3* and *clv1-101 crn elf3* with all single mutant control alleles. **c**, *CLV*-mediated FM stem cell regulation is sensitized to auxin levels. Carpels per flower at 23°C for Col-0, *yuc4*, *clv1-101*, *clv1-101 yuc4*, *clv3-9*, *clv3- 9 yuc4*. 10 carpels counted per plant, n=8-24. Statistical comparisons based on Mann-Whitney test (**a**) where * indicate p-value < 0.0001 and ns indicates not significant. Statistical groupings based on significant differences using a Kruskal-Wallis and Dunn’s multiple comparison test correction **(b, c**). Significance is defined as p-value < 0.05 and “+” indicates mean (**a-c**).

**Extended Data Fig. 5:**
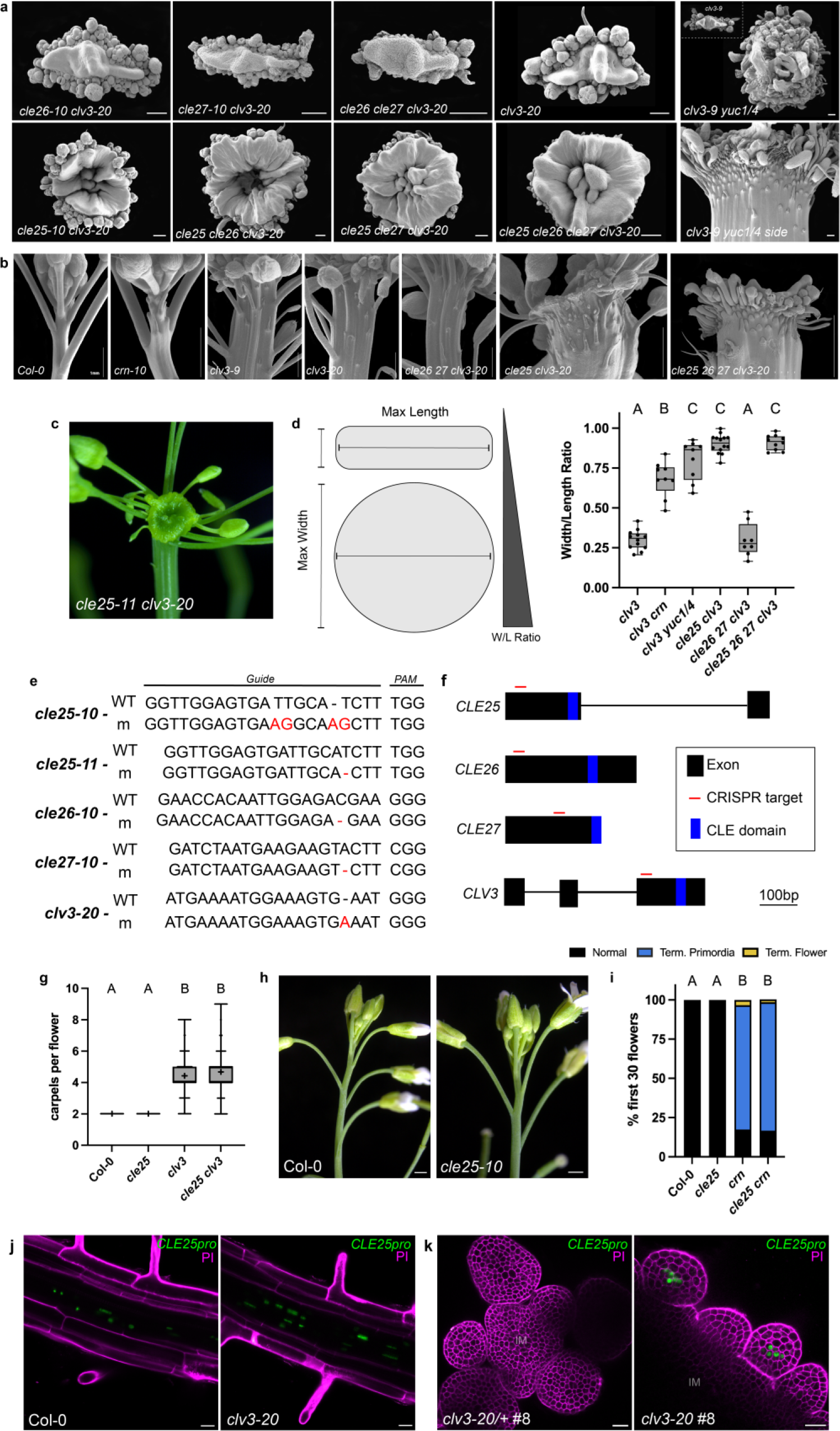
*CLV3* acts redundantly with *CLE25* and auxin in IM function. **a-c**, *cle25* and *yuc1/4* mutations enhance *clv3* IM phenotypes. **a,** SEM images of IMs of *cle26-10 clv3-20*, *cle27-10 clv3-20*, *cle26-10 cle27-10 clv3-20*, *clv3-20, cle25-10 clv3-20*, *cle25-10 cle26- 10 clv3-20*, *cle25-10 cle27-10 clv3-20*, and *cle25-10 cle26-10 cle27-10 clv3-20*. *clv3-9 yuc14* and *clv3-9* inset at same magnification for size comparison. *clv3-9 yuc1/4* side view. **b**, Flower primordia termination visualized with SEM side view of IMs in Col-0, *crn, clv3-9, clv3-20, cle26- 10 cle27-10 clv3-20, cle25-10 clv3-20, cle25-10 cle26-10 cle27-10 clv3-20.* Scale bars 1mm. **c**, Inflorescence of the *cle25-11* allele with *clv3-20* mutation showing characteristic disk-like IM phenotype. **d**, Quantification of disk meristem expansion using ratio of maximum width and maximum length of *clv3-20, clv3-20 crn, clv3-9 yuc1/4, cle25-10 clv3-20, cle26-10 cle27-10 clv3- 20 and cle25-10 cle26-10 cle27-10 clv3-20.* **e**, CRISPR guide and PAM sites for *CLE* genes indicating SNPs and indels in red or with dash. WT, wild-type; m, mutant. **f**, Gene diagrams of *CLE* ORFs with red line indicating CRISPR target positioned before the CLE domain indicated by the blue rectangle. **g**, carpels per flower of Col-0, *cle25-10, clv3-20* and *cle25-10 clv3-20* (n=12). **h-i**, *cle25* alone does not enhance *crn*. **h**, Inflorescence images and **i**, quantification of termination of Col-0, *cle25-10, crn* and *cle25-10 crn* (n=9). *CLE25* is expressed in root phloem cells independent of *clv3*. **j**, Confocal images of *CLE25pro::YPET-H2AX* (green) of Col-0 and *clv3-20* in roots. **f**, *CLE25* is ectopically expressed in *clv3-20* floral primordia. Col-0 (n=4) and *clv3-20* (n=4) with same single transgene *CLE25pro::YPET-H2AX* (green) insertion line (#8) in IM and FMs (PI, magenta). Scale bar 20µm (**e, f**). Statistical groupings based on significant differences using a Kruskal-Wallis and Dunn’s multiple comparison test correction (**d, g, i**) where significance is defined as p-value < 0.05.

**Extended Data Fig. 6:**
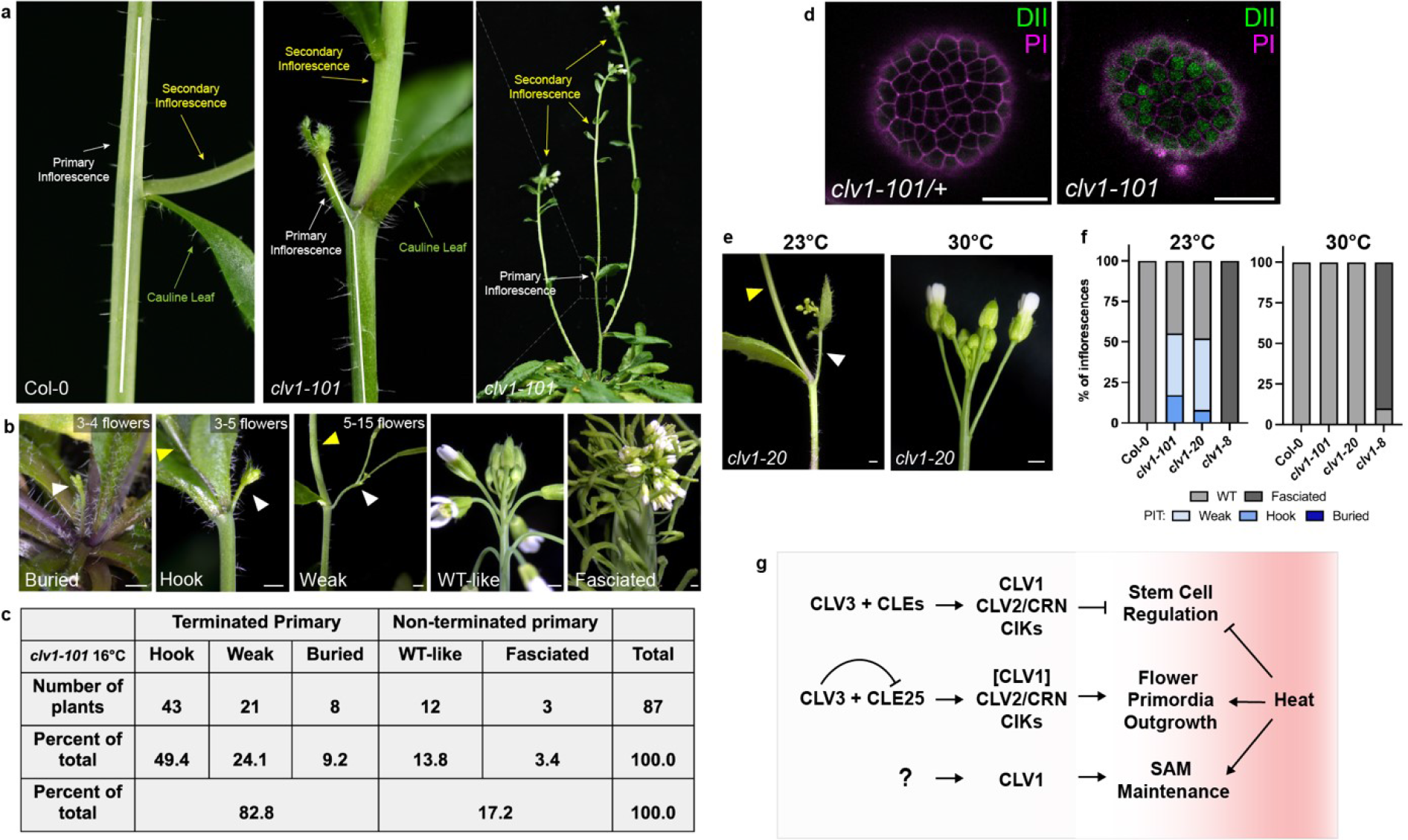
CLV1 is required for IM maintenance. **a-e**, *CLV1* is required for primary IM maintenance. **a,** Side view of primary inflorescence with arrows pointing to primary inflorescence (white) with cauline leaf (green) subtending the secondary inflorescence (yellow) in both Col-0 and *clv1-101* revealing primary inflorescence termination (PIT) in *clv1-101*. **b**, Categories: buried, hook, weak, WT-like, and fasciated. Range of flowers that emerge in each category before PIT. **c**, Population analysis of *clv1-101* at 16°C (n=87). **d**, *UBQ10::DII-Venus* in L1 cells of the IM in *clv1-101/+* and *clv1-101* homozygous mutant undergoing PIT. The same transgenic insertion line was used in d. 4/4 WT *UBQ10::DII-Venus* and 4/5 *clv1-101 UBQ10::DII-Venus* (green) independent transgenic lines showed representative expression. *clv1-20* allele inflorescence undergoing PIT at 23°C but not at 30°C (PI, magenta) (**e**) and quantification of PIT in Col-0, *clv1-101, clv1-20* and *clv1-8* plants (**f**, n=15). **g**, Summary of interactions between CLEp signaling and heat on auxin dependent stem cell regulation, flower primordia outgrowth, and SAM maintenance. Scale bars 10mm (**b, d, e**) and 20µm (d).

